# Emergence of flagella-like oscillations in single microtubules driven by collective dynein transport

**DOI:** 10.1101/2022.09.11.507451

**Authors:** Shivani A. Yadav, Neha Khetan, Dhruv Khatri, Chaitanya A. Athale

## Abstract

Flagellar and ciliary oscillations result from a combination of stereotypical axonemal geometry, collective mechanics of motors, microtubules (MTs), elastic linkers and biochemical regulation. However, the minimal essential components and constraints resulting in flagellar oscillations remain unclear. Here, we demonstrate that periodic, low-frequency waves of flagella-like oscillations *in vitro* emerge from a ATP-driven collective molecular motor transport of MTs clamped at one end. The spontaneous oscillations arise without any external forcing and can be explained by an in *silico* model of molecular motor binding driven MT bending and buckling followed by motor detachment driven ‘recovery’ stroke. We demonstrate that transitions in single MT patterns between flapping, flagellar-beating and looping are determined solely by the self-organization of collective motor transport and filament elasticity.

## Introduction

Cilia and flagella are eukaryotic cellular appendages found in diverse phyla that are responsible for motility, sensing and generating extracellular flows. Their core is referred to as the axoneme and consists of a cylindrical arrangement of typically 9+2 microtubule (MT) doublets with minus-end directed motors, dyneins, linking these doublets along the circumference (*1*). The elastic nexin connect the central doublet to the outer ones constraining them like spokes (*2*). When dynein motors step on adjacent doublets, the constraints of intact flagella result in propagation of bending forces along the length (*3*). Flagellar beating is thought to be self organized and diverse species-dependent patterns are thought to emerge from the detailed mechanics of the molecular components (*4*–*6*). The nature of the flagellar waveform was demonstrated to depend on the specific nature of load-dependent detachment of dynein and the immobilization of the axoneme at the base using bull sperm flagella in both theory and experiment (*7*). At the same time distinctions between waveforms have been made classifying them as ciliary if the curvature originates from the base and propagates towards the tip, and flagellar for tip-to-base propagation (*8*). Flagellar beating can however also reverse direction (base-to-tip) seen in random, transient switching as well as that induced by extracellular viscosity and Ca^2+^ion concentration (*9*, *10*). Filament length also appears to play a role as seen with beat wavelength dependence seen in *Chlamydomonas* flagella (*11*). Beat initiation was hypothesized to be a self organized radial sequence of motors switching by activation and de-activation, described by the ‘geometric clutch model’ (*12*). A minimal 2D model that extends this hypothesis, has been shown capable of reproducing the emergence of planar beating waves (*13*). In *vitro* sliding of demembranated axonemes demonstrated cooperative motor transport since sliding velocities were higher than single molecule velocities, but the approach did not produce the curvature seen in intact axonemes suggesting the absence of constraints (*14*). Propagation of curvature was reported previously in actin transport by myosin sheets in a ‘gliding assay’ resulting in spirals that were modeled as a result of filament elasticity and motor activity (*15*). Transient waveforms of bending have also been reported for free MTs bound to clusters of the plus-end directed motor kinesin that are transiently surface-bound (*16*). These are generally considered to arise due to surface defects and do not lend themselves to a controlled study. While MT bundles driven by complexes of kinesin and crowdants were shown to produce synchronized base-to-tip ciliary waves *in vitro* (*17*), the study lacked control over MT length and bundle density. This shortcoming was also common to a demonstration of MT length and motor density control of cilia by reconstitution (*18*) and the bending wave generation seen in axonemal dynein driven demembranated axonemal MT sliding (*19*). Therefore an experimental setup where the components and constraints are clearly defined, could help to better understand the principles underlying flagella-like MT beating. This would also allow a comparison to theoretical models of molecular motor mechanochemistry and cytoskeletal elasticity.

Here, we show flagella-like oscillations that emerge from plus-end clamped single MT filaments whose free ends are subject to forces from immobilized dynein motors acting along their length in a ‘gliding assay’ format. We have used the minimal cytoplasmic yeast dynein as a force-generator, due to its emergence as a model for minus-ended motors with detailed single molecule mechanics (*20*), asymmetry in force dependent detachment (*21*, *22*) and collective transport (*23*, *24*). Our minimal system results in cyclical wave-like beating, similar to flagella. In parallel, we have developed a computational model using discrete stochastic interactions between motors and microtubules. The model predicts the limits within which regular oscillations of the filaments emerge with motor density, MT length and proportion of the MT length pinned to the surface.

### Flagella-like beating of single MTs in dynein collective transport with plus-end pinning

Our approach involves only two components: MTs with biotinylated plus ends immobilized on the surface by streptavidin and nanobody immobilized cytoplasmic dynein motors layered on the same coverslip surface that bind along the length of MTs. We have used the minimal yeast cytoplasmic dynein (*20*) that has become a model dynein due to ease of purification and high processivity in driving unidirectional filament transport in ‘gliding assays’ (*23*). The MT geometry with plus-ends clamped and free minus-ends that can bind to motors, in presence of ATP results in the collective force generation by dyneins. This results in a bending force generated akin to a cantilever clamped at one end while experiencing compressive force along its entire length due to motors acting to transport the filament in the direction of the clamp (plus-end), resulting in increasingly curved structures (Figure 1(a)). The biotin and Alexa Fluor-488 tagged plus-ends (green) anchored by streptavidin on the glass surface remains straight, while the rhodamine-tubulin tagged free-ends bend and buckle, producing increasing elastic force. Once the filament bending exceeds the energy of the MT-motor attachment, the motors detach resulting in a ‘recovery’ stroke, mimicking flagellar and ciliary dynamics (Figure 1(a), Video SV1). This process takes ~400 s for one cycle to complete and the time-projected contours of the filament emphasize the oscillatory nature of the filament movement (Figure 1(b)). The movement of the free minus-end of the filament is seen in the dynamics of the X- and Y-coordinates and the elevation angle of the free tip (Figure 1(c)). The propagation of filament curvature from base to tip is illustrated in the kymograph of the tangent angle along the filament contour with time, where MT free ends appear to complete one wave in ~300 s, corresponding to a frequency of 3 mHz (Figure 1(d)). We also find filaments assembled with biotinylated plus-ends and rhodamine-tubulin to visualize both bound and free parts also show similar patterns (as compared to dual labelled filaments) with the tip oscillating with ~5 mHz frequency (Figure S1, Video SV2). Multiple fields of view demonstrate that not all filaments get captured by the streptavidin bonds, and some filaments are transported in a processive manner of a ‘gliding assay’ (Video SV3). We find an optimal density of streptavidin and nanobody is required to achieve the flagella-like patterns, as a result of which some filaments may not encounter a pinning site at all, and instead simply be transported. While comparable *in vitro* approaches have been taken in the past, our approach is distinct due to differences in terms of either the motor type used, geometry of the system or the regular oscillations. For example when kinesin was used to drive MT bending due to clamped minus-ends (*25*), the experiments did not result in oscillations. A similar mechanism was invoked to explain in vivo observations of highly curved MTs (*26*). The absence of wave-like oscillations can be explained by both the differences in the motor mechanics of force generated, speed and detachment force as well as conditions such as motor density and geometry of the force exerted. On the other hand when oscillatory beating patterns have been observed previously, they were based on non-specific pinning of kinesin-complexes and accidental clamping of MTs (*16*). In the case of bundles of MTs beating driven by kinesin-complexes the number and lengths of MTs were unclear (*27*). Therefore our approach demonstrates both a simpler set of components and constraints, that result in the emergence of flagella-like patterns in single MTs.

**Figure 1:**
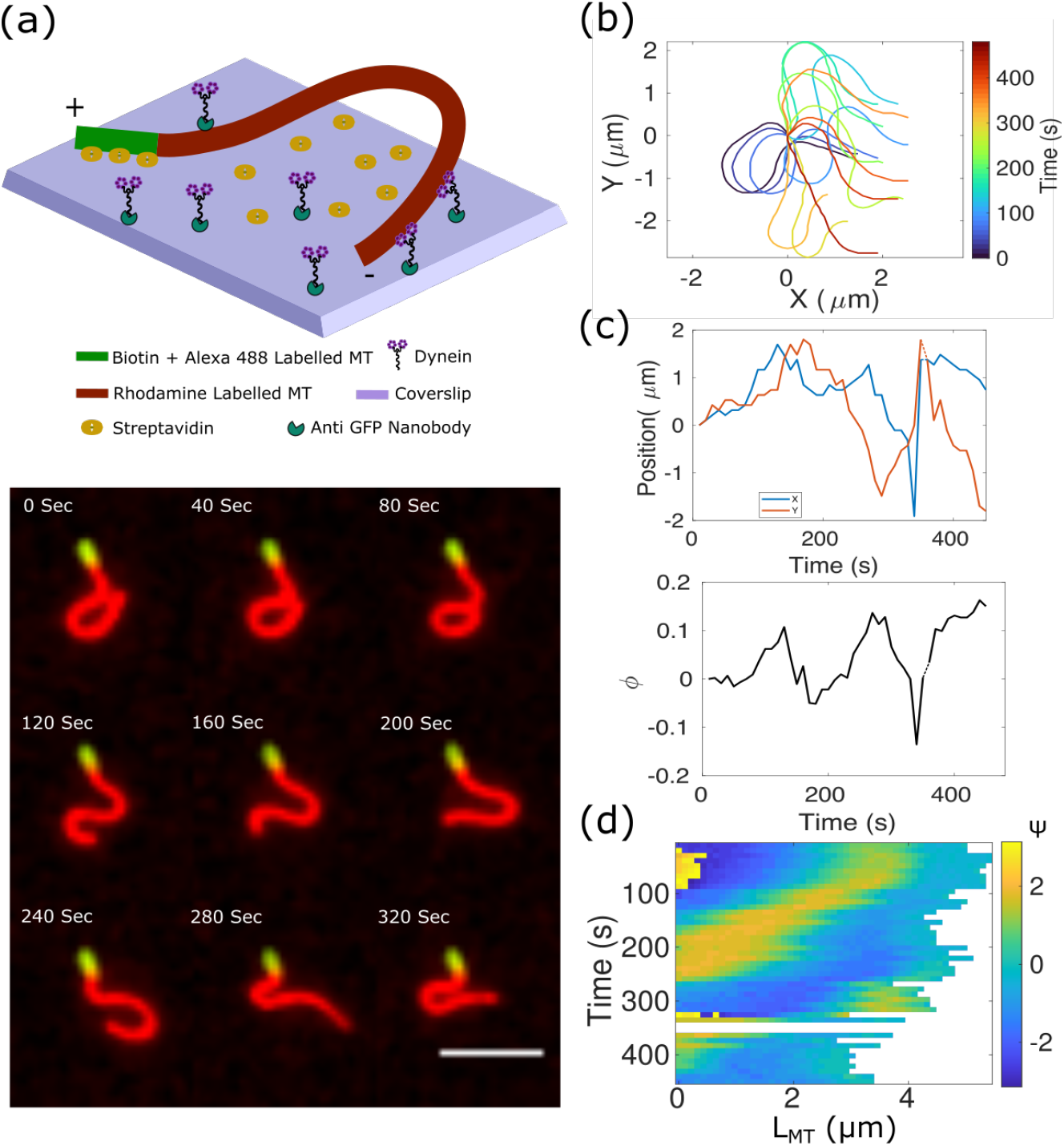
Dynein driven flagella-like MT beating reconstituted *in vitro* with biotin-based pinning. (a) The emergence of flagella-like beating of an MT pinned to the surface by the plus-end, with the minus-end oscillations driven by surface-immobilized dynein as seen in the (top) schematic and (bottom) representative fluorescence microscopy time-series. In experiment the plus- and minus-ends were labelled using Alexa-488 (green) and rhodamine (red) tubulin respectively (Materials and methods). Plus ends also had biotinylated tubulin that bound to streptavidin coated on the glass surface. Time in seconds. (b) Filament contours from experiment were projected in time at an interval of 30 s over the entire trajectory (color-bar: time in seconds). (c) The minus end of the MT that is free is analyzed for the position in X (blue) and Y (red) as a function of time (top), and its angular elevation *ϕ* (bottom) was plotted. (d) The local tangent angle along the MT contour, *ψ*, is plotted with time as a kymograph at 10 s intervals. Colors represent the angle in radians. Scale bar: 4 *μ*m.

Our observation of single filament beating is reminiscent of ciliary waveforms propagating from base to tip (*8*), despite the absence of the complex machinery and geometry that drives bonafide ciliary and flagellar dynamics. At the same time the complex curvature compared to the almost planar waves of cilia, makes it more similar in form to flagella. While previous reports of in *vitro* reconstitution of MT-motor interactions in a ‘gliding assay’ format have shown a range of patterns from MT buckling of biotin-pinned MTs with a low kinesin density (*25*), waving and rotational motion (*15*, *28*) and loop formation at high motor density (*29*), such periodic waves were not reported. The generality of such structures was illustrated by spirals and flagellum like bending of actin formed when filament tips were pinned to surface defects and myosin motors generated forces (*15*). The wide-spread observation of patterns of cytoskeletal filaments driven by motors-MT-Kinesin, MT-Dynein and actin-myosin-suggests flagella-like oscillations are general patterns requiring a minimal set of components. We therefore proceeded to develop a computational model to examine the principles governing the patterns, and explore the sensitivity of the system to motor density and MT length.

### Model of pinned MTs and dynein transport recapitulates flagella-like oscillations

Can a computer simulation of just two components-MTs and motors - reproduce the regular beating patterns seen in experiment? To address this, we configured a model using an agent-based stochastic simulator Cytosim with (i) MTs modelled semi-flexible polymers with plus-ends clamped to the surface and minus-ends free and (ii) minus-end directed molecular motors that have a linear force-velocity relation, probabilistic binding and force-dependent unbinding rate and stochastic stepping (Figure 2(a)). Equations governing the model are described in detail in the model description (Supplementary Material) while parameters are taken from literature where possible (Table S1). We model the clamping of MT-plus ends by biotin-streptavidin linkage by assuming the plus-end of the MT is attached to the surface by a stiff spring (spring constant *k_a_* of 10^3^ pN/μm) that does not detach. Simulations are initialized with a straight geometry of the free-end of the MTs, and these on binding to motors with the plus-end clamped spontaneously result in the emergence of periodic beating patterns (Figure 2(a), Video SV4). The patterns of bending appear symmetric between the initiation and recovery strokes in time-projected filament plots (Figure 2(b)). The periodic movement of the free tips with some stochasticity can be observed in the X- and Y-position (Figure 2(c)) and angular elevation (*ϕ*) plots (Figure 2(d)). One cycle completed in ~300 s corresponding to a frequency of ~3 mHz, corresponding closely to experimental observations. It is noteworthy that these oscillations are slower than intact flagella and cilia that have a frequency ranging between 1 and 100 Hz (*32*). The base to tip propagation of curvature is seen in the kymograph of the local curvature of the filament (Figure 2(e)) measured by the tangent along the contour (*ψ*, Equation 4) is qualitatively similar to experiment. Thus without any optimization of parameters we find our simulations are qualitatively comparable to experiments in terms of base-to-tip bending, ‘recovery’ stroke, amplitudes and time scales.

**Figure 2:**
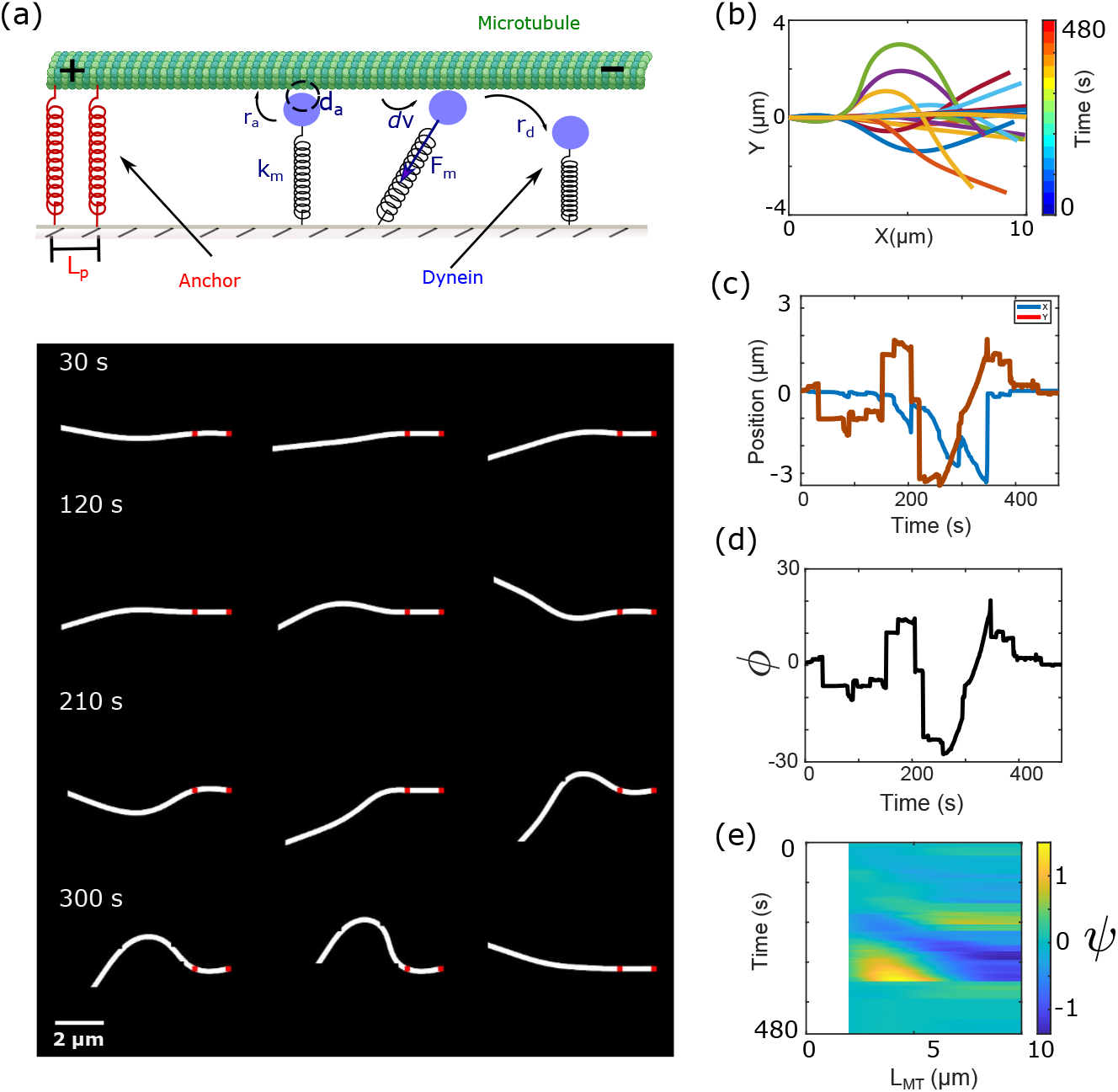
Simulating motor driven flagellar beating of a pinned MT. (a) *Top:* MTs are modeled to interact with surface immobilized dynein motors (blue) and anchors (red) that pin them at their plus ends over the length of (*L_p_*). Motor heads (blue circle) stochastically attach to a MT within a distance *d_a_* at the rate *r_a_*, take steps *d_v_* and can detach at a force-dependent rate *r_d_*. The motor is treated as a spring with a constant *k_m_* resulting in a force *F_m_* when extended. *Bottom:* A representative image time-series of the 2D simulation of filament beating. Time in seconds and scale bar: 2 *μ*m. (b) Filament contours are projected in time at an interval of 30 s during a 480 s period of simulation. The free minus end of the MT is analyzed for (c) the position in X (blue) and Y (red) as a function of time and (d) its angular elevation, *ϕ*. (e) The local tangent angle along the MT filament *ψ*, is represented as a kymograph (X: length of the MT, Y: time). Colors denote the angle in radians. Here, filament length L_*MT*_ = 10 *μ*m, motor density *ρ_m_* = 10 dyneins/*μ*m^2^, pinned MT length L_*p*_ = 2 *μ*m

### Motor number dependent transition from MT beats to loops

A previously developed general model of spirals of actin driven by myosin collective transport predicted a scaling relation of the radius of the spiral *R* ∼ (*k_B_Tξ_p_/f*)^1/3^ where f is the linear force density and *ξ_p_* the persistence length (*15*). While MTs are more rigid (*ξ_p_* ~ 1 mm) compared to actin (*ξ_p_* ~ 10 μm) it remained to be seen if spiral like patterns emerged in our system. Experimentally, MT length and motor density are the easiest parameters to vary.

We proceeded by varying MT filament lengths (*L_MT_*) over a range of 5 to 25 μm based on typical experimental observations. At the same time the density of dynein motors (*ρ_m_*) was varied between 0 and 10^3^ motors/*μ*m^2^, based on previously reported experimental values (*23*, *30*). The time-projected images of filament contours demonstrate the formation of multiple, distinct patterns, which we broadly classify as (*a*) fluctuations and bending, (*b*) flagella-like beating and (*c*) looping (Figure 3(a)). Fluctuations are observed in the complete absence of motors, similar to those reported for cantilevers in a thermal bath. As the motor density increases, the patterns appear to depend on the MT length. Short filaments of length 5 *μ*m subjected to increasing motor activity appear to undergo bending to one side, do not recover as seen in a representative plot with *ρ_m_* = 10^1^ motors/μm^2^. At high densities filaments buckle and undergo *looping*, when *ρ_m_* = 10^3^ motors/μm^2^ (Video SV5). This loop structure emerges at lower motor densities for longer filaments (10 to 25 μm). At the same time along the diagonal of the *ρ_m_* and *L_MT_* axis, we observe a narrow region of patterns that resemble *flagella-like beating*. Representative kymographs of motor density dependence for 15 *μ*m long MTs show regular beating only when *ρ_m_* = 10 motors/*μ*m^2^ (Figure 3(b)), while at a constant ‘ideal’ density *ρ_m_*=10 motors/μm^2^ periodic beating emerges when MT lengths of ≥ 15 *μ*m (Figure 3(c)). These results suggest the onset of regular beating requires a minimal MT length, while for sufficiently long MTs, the motor density (*ρ_m_*) determines the beating frequency.

**Figure 3:**
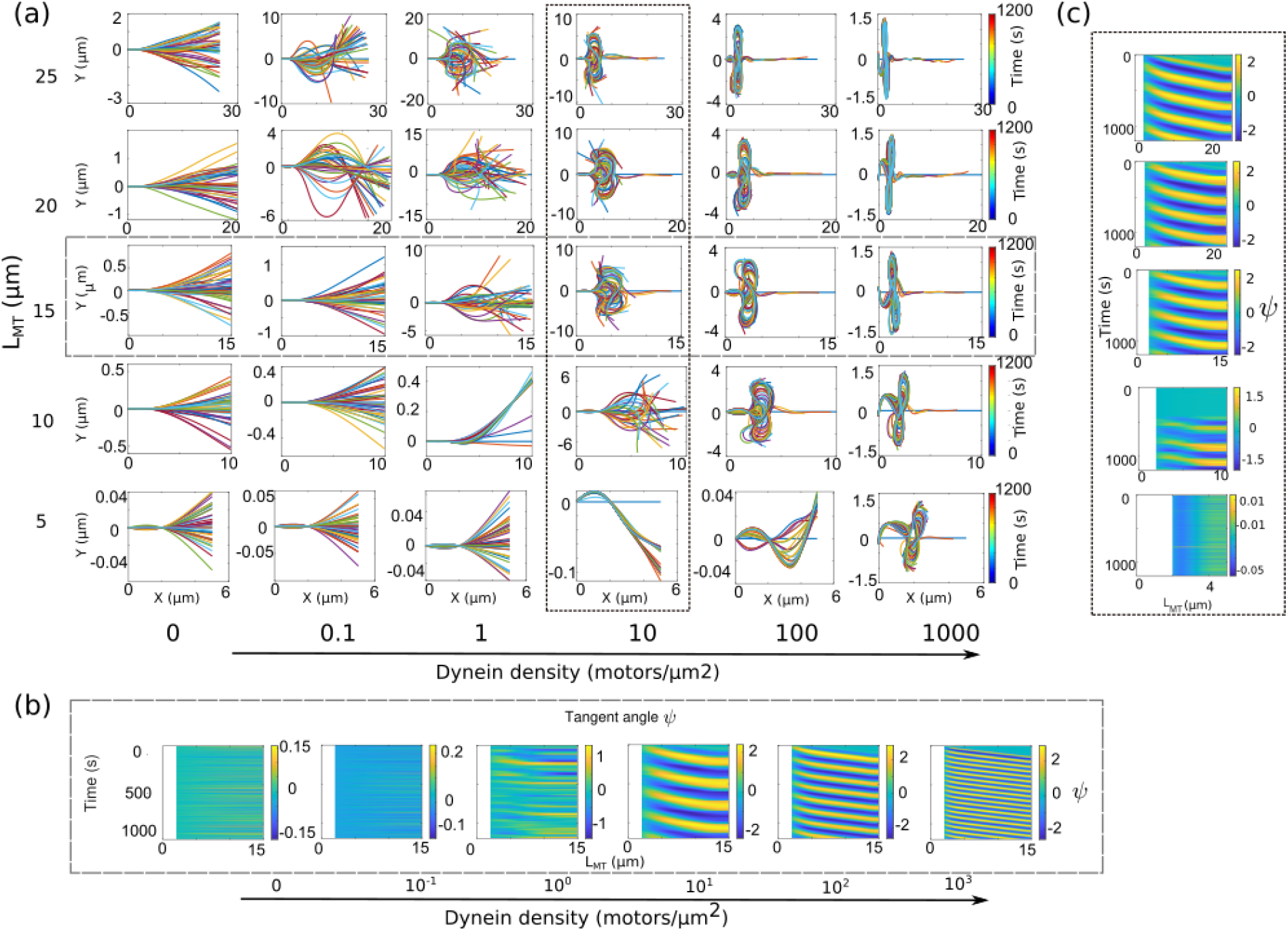
Simulating the effect of MT length and motor density on flagellar beating. (a) The dynamics of filaments are simulated with MT lengths ranging from 5 to 25 *μ*m (rows) and motor densities ranging from 0.1 to 10^3^ motors/*μ*m^2^ (columns). The projected filament contours are plotted at 30 s intervals over a total time of 1200 s. (b,c) Representative kymographs of the local tangent angles (*ψ*) plotted as a function of position along the filament (x-axis) and time (y-axis) are plotted. Dashed lines indicate either that (b) the motor density was increased while keeping MTs lengths at 15 *μ*m or (c) MT length was increased, while motor density was 10 motors/*μ*m^2^.

In simulations filament deformation originates from the pinned-end (Figure 3(b,c)). This is likely to be due to the polarity of the microtubule-motor system, and the nature of the cooperative force generation of dynein (*31*). Although such base-to-tip propagation is considered to be ciliary (*8*, *13*), our observation of complex curvatures with more than one principal curvature and occasional out of plane movement suggests more flagella-like behavior. The ‘looping’ observed for increasing dynein densities suggests a qualitative change due to the higher force density, resulting in higher buckling modes at shorter length scales. Taken together, we find MT beating is predicted to arise at a critical motor density that determines the amount of force generated and MT length, that determines both the buckling force and elastic force resulting in motor detachment and ‘recovery’. We proceeded to test whether the relative proportion of the filament length that was immobilized could also play a role.

### Proportion of pinned and free MT length affects the beating patterns

Our simulations predict that sustained MT beating results from a narrow range of motor density and MT lengths (Figure 3), with ≈10% of the filament length pinned to the surface. From linear elasticity theory, we expect that the energy required to bend a filament depends on the length, *E_bend_* ∝ *L*. This would suggest that if we varied the proportion of the free-end of the filament as a fraction of the total length, we should alter the nature of patterns observed. We observe flagellar beating emerges in simulations when the length pinned is 0.5 μm and 1 μm, i.e. 5 to 10% of the total filament length from a scan over a wider range (Figure 4(a)). The fraction of pinned length determines the emergence of significant filament bending. When the proportion of pinned filament is small i.e. 1%, *L_p_* = 0.1 μm, the filament appears to rapidly oscillate between looped forms. For a very large pinned fraction, 20%, *L_p_* = 2 μm, it only results in minimal MT bending that are transient (Figure 4(b)). All this suggests that the linear force density (force per unit length) is likely to play a role. To test this, we simulated a ten-fold higher motor density and found filament ‘swiveling’ when the fraction pinned was 1% of total length (*L_p_* = 0.1), while spirals and loops formed when *L_p_* = 2 μm (20%) (Figure S2). We also find for a large fraction of the filament length being pinned, as in the case of MTs with 20% of the length pinned (*L_p_* = 2 μm), filaments show variability between simulation runs ranging from continuous beating in some cases, with others showing transient beats interspersed with pauses. In experiment, we find that filaments with increasing pinned proportion-3.6%, 6.4% and 22.4% of the total MT length-transitioned from looped form to periodic beating (Figure 4(c)). The tip position dynamics of the representative experiments suggest a transition from slow to regular oscillations that eventually disappear altogether with peak-peak frequency of 2-5 mHz (Figure 4(d)).

**Figure 4:**
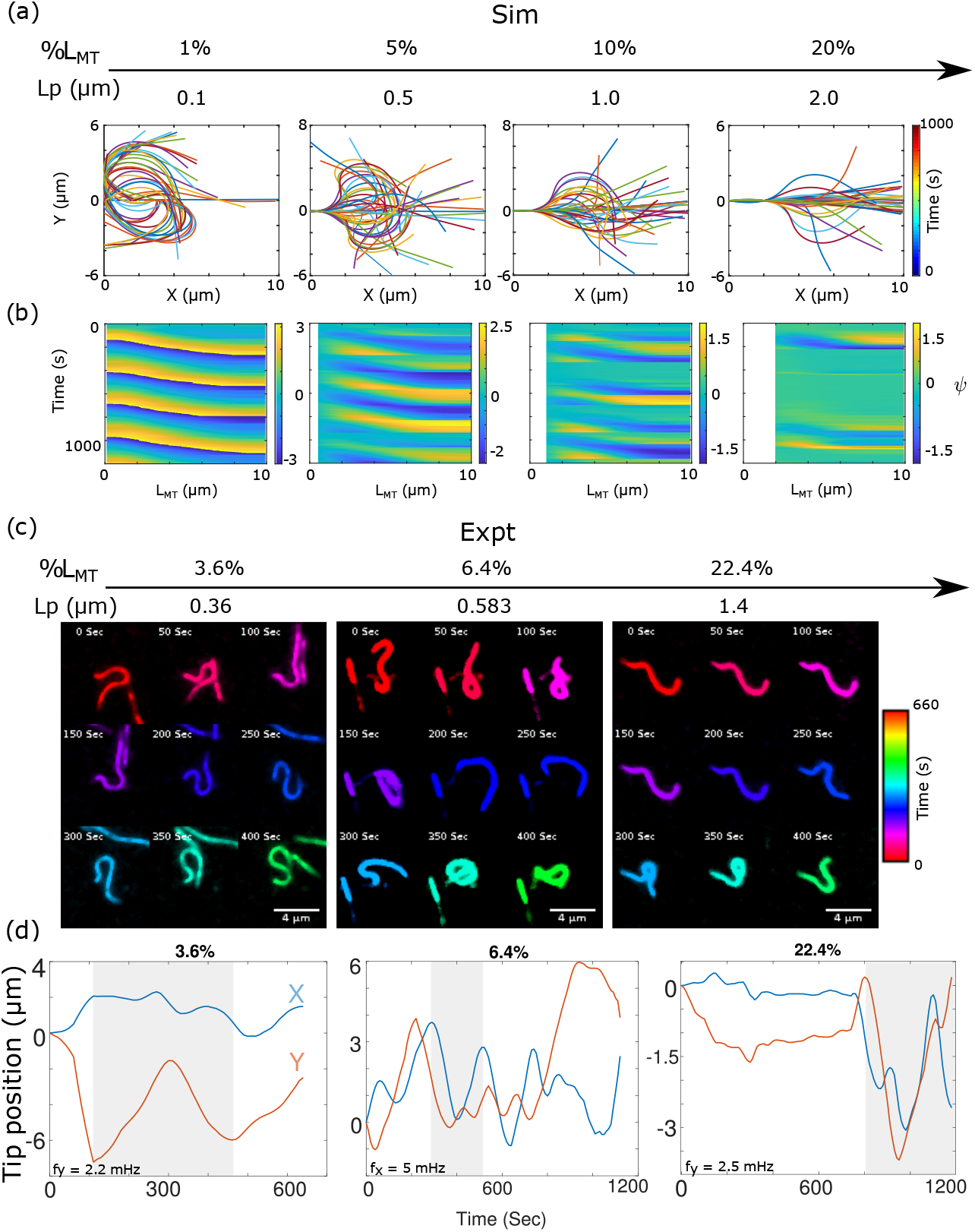
Proportion of MT length pinned affects MT-beating patterns in simulation and experiment. (a) Representative simulations of 10 *μ*m long MTs and constant motor density (*ρ_m_* = 10 motors/*μ*m^2^) with increasing pinned length (Lp) 0.1, 0.5, 1 and 2 *μ*m (left to right) are visualized as time projected contours at 30 s intervals with corresponding (b) kymographs of the tangent angles (*ψ*) along the contours. (c) Experimental microscopy data is represented as a montage of time for filaments of increasing proportion of the length pinned - 3.6%, 6.4% and 22.4% (corresponding to Video SV6(a-c)).Colors indicate time frames (time interval 50 s). (d) X and Y coordinates of the beating MTs as a function of time for MTs with increasing proportion of the length pinned (data smoothed for clarity) is used to infer the frequency using X (*f_x_*) or Y (*f_y_*) coordinates depending on the regularity of the profile. Grey areas highlight one cycle of oscillations.

Taken together our simulation predictions of the proportion of the pinned MT length are confirmed in experiment, and suggest that the free-filament length and motor density are critical parameters that determine the emergence of flagella-like beating patterns.

### Phases of flagellar patterns driven by MT length and motor density

Our experiments and simulations demonstrate the systematic emergence of periodic beating patterns arising from a simple 2D system that are specific to the relative proportion of the free filament and motor density. A wider scan of lengths and densities (Figure S3) predicts that short filaments (~ 5 μm) will only undergo (I) *fluctuations* even when *ρ_m_* is varied from 0 to 10^2^ motors/μm^2^, while long filaments (~ 20 *μ*m) will exhibit flagella-like (II) *beating* motion even at lower dynein densities of 10^-1^ dynein/*μ*m^2^, while filaments undergo (III) *looping* when the MTs are long and motor density sufficiently high (Figure S4(a),(b)). In order to objectively capture the shape of the filament and distinguish between the different qualitative forms, we develop a measure that is the logarithm of the ratio of maximal distance explored along x- and y-axes (Equation 5), and find a clear separation (Figure 5). The metric *S* defines the transition to the qualitatively distinct states as summarized here:

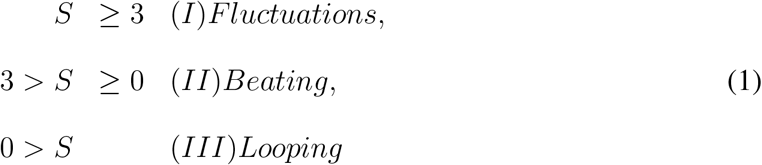

**Figure 5:**
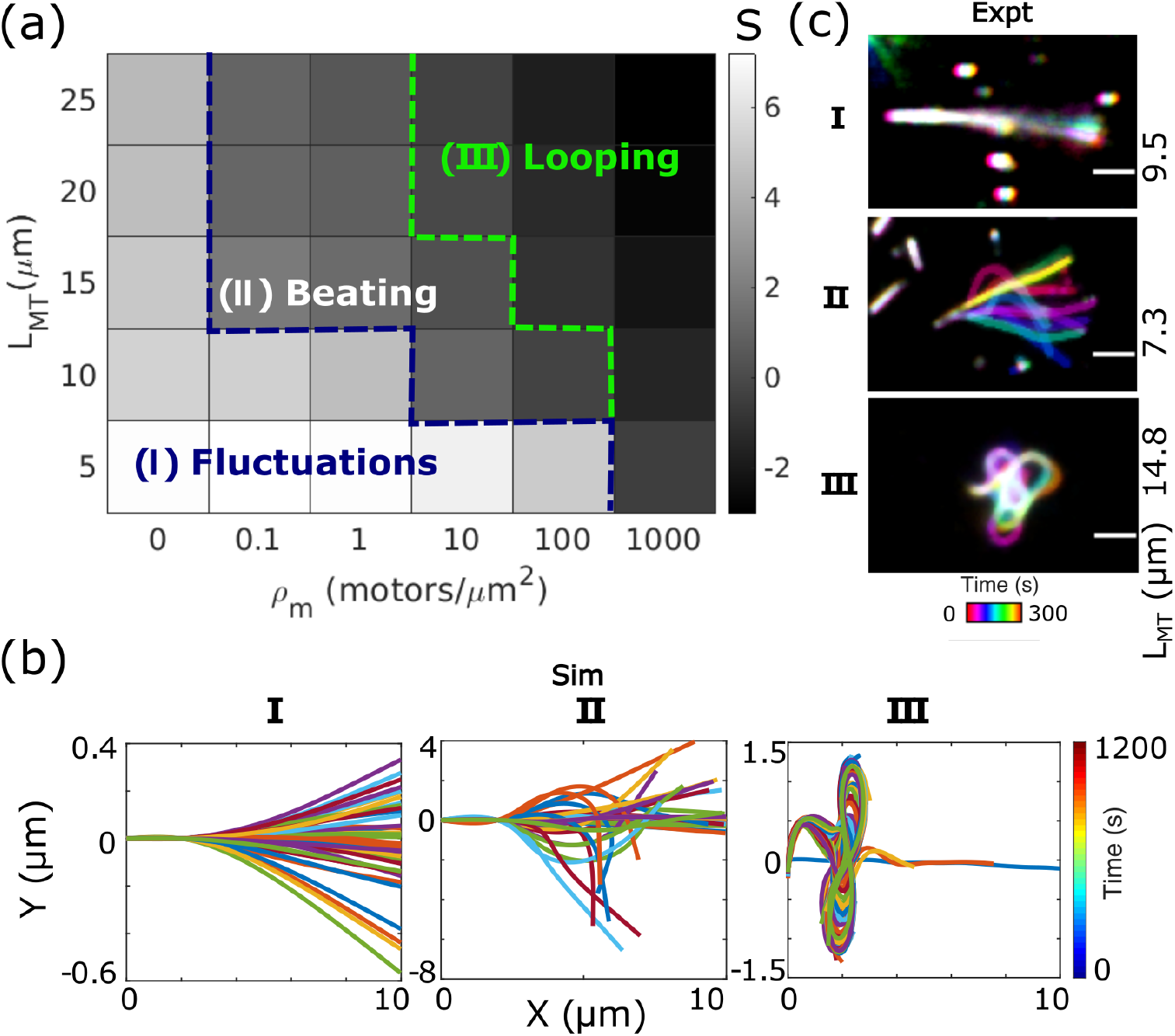
Phase diagram of qualitative differences in beating with motor density and MT length. (a) The parameter space for varying motor densities (*ρ_m_*) and MT lengths (*L_MT_*) are divided into three distinct regions based on the derived measure, 〈*S*〉 that serves as a measure of qualitative differences in filament dynamics (averaged over 10 runs for each parameter). The three regions correspond to (I) fluctuations, (II) beating, and (III) looping. (b, c) Representative images of time-encoded filament contours for each type of motion are compared between (b) simulations and (c) experiments. (b) The representative simulated MTs are 10 *μ*m long with 2 *μ*m pinned, while the density of dynein was increased: (I) 0.1, (II) 1 and (III) 10 motors/*μ*m^2^. (c) Representative experimental data of pinned MTs (I) in the absence of motors (*L_MT_* = 9.5 *μ*m) and with (II, IIII) a motor density of 46 dynein dimers/*μ*m^2^ for MT lengths (II) 7.3 *μ*m and (III) 14.8 *μ*m. Color-bar: time (s). Scale bar: 2 *μ*m.

In the exceptional case of single pivot points of filaments in the absence of motors, this variable S becomes 0, a signature of diffusive swiveling. Experimental data matches the patterns with the absence of motors leading to thermal oscillations (I), while the presence of a motor density of 46 dynein dimers per *μ*m^2^ with 7.3 *μ*m MT results in flagella like oscillations (II), while a filament of 14.8 *μ*m length at the same motor density exhibits looping (Figure 5(c)). Further, we observe these distinct MT patterns can emerge for comparable MT lengths and dynein density under varying proportion of pinned lengths (Figure S5), suggesting that motor numbers, MT length and the MT pinning can determine the nature of MT patterns.

The abrupt transition between fluctuating and flagellar beating states merits further exploration of potential phase transition like behavior. The flagellar beating frequency (phase II) is ~0.02 Hz at the mid-length of the MT (Figure S4). This is lower by multiple order of magnitudes as compared to the frequency of intact cilia and flagella whose frequencies range between 1 and 100 Hz (*32*). Part of the explanation for the difference is likely to be the much simpler nature of both our experimental and simulation setup, whereas intact flagella have more complex geometries experience higher magnitudes of forces. Additional factors could include the differences in motor mechanics, since for example yeast cytoplasmic dynein *in vitro* collective velocity is ~ 0.1 *μ*m/s (*20*, *23*) while the outer arm dynein (OAD) from *Tetrahymena* transports MTs in a ‘gliding assay’ at a velocity of 5 *μ*m/s (*33*). In addition detailed mechanics of these motors are also likely to differ. Some of these features can in future be tested both in experiment and simulation.

In summary we define the minimal experimental constraints by which a single filament clamped by an end to a surface, together with minus-end directed dynein motors can spontaneously result in regular, oscillatory flagella-like beating patterns. A computational model of semi-flexible filaments and simple and spring-like motors with stochastic binding and force dependent unbinding, can reproduce the observed behavior. The simulations also predict a minimal motor density, ideal MT length and a the proportion of MT pinned, thus defining the constraints. Our simulations were validated by comparison to experiments. While flagella and cilia in vivo have a stereotypical 9+2 geometry of doublets with crosslinkers and spokes, this minimal system suggests that the elastic nature of end-clamped MTs and driven to bend can produce qualitatively similar patterns as observed in bonafide flagella. The almost ≈10^3^ lower frequency could originate from multiple sources-the higher rigidity, more complex geometry, nexin linkers, spokes and potentially greater number of motors. Indeed in simulations we find very little effect of single-motor speed on beating frequency, suggesting the geometry of the axoneme is critical for the observed high frequency oscillations. At the same time the density and length dependence suggests that qualitative transitions in flagellar oscillations are to be expected from not just the minimal but even potentially the complex flagellum. The use of the cytoplasmic dynein suggests that the force produced by an unrelated cytoplasmic motor is sufficient to overcome the elastic energy of bending of an MT. In some senses the biotin-streptavidin pinning site acts as an implicit constraint that in the axoneme is provided by the nexin cross-links along the circumference of the axoneme. To that extent in future a filament doublet or higher order complexity could be used to test the validity of this assumption. The self-organized nature of the observed patterns suggest that while layers of regulation and coordination are reported to be involved in flagellar and ciliary oscillations, the mechanics alone are potentially a modular element which is further constrained for more robust and faster beating. These insights taken together can be of use in designing a minimal synthetic flagellum in future.

## Acknowledgments

We thank Kheya Sengupta for discussions.

## Funding

SAY is supported by a senior research fellowship by CSIR-India (09/936(0261)/2019-EMR-I). NK was supported by a postdoctoral fellowship by the Indo-French Centre for Promotion of Advanced Research (CEFIPRA) 62T5-D. DK is supported by a fellowship from the Dept. of Biotechnology (DBT/JRF/BET-18/I/2018/AL/188). This work was funded in part by CEFIPRA grant 62T5-D to CAA.

## Author Contributions

SAY performed the experiments, NK performed simulations and wrote the analysis codes, DK performed the image-analysis, SAY, NK and DK prepared the figures, NK, SAY and CAA wrote the manuscript, CAA conceptualized and supervised the work.

## Competing interests

The authors declare they have no competing interests.

## Supplementary materials

- Model
- Materials and methods
- Tables S1
- Figures S2 to S6
- Videos SV1 to SV6

### Model

#### Microtubules

Microtubules (MTs) are modeled as semi-flexible filaments. All filament lengths (*L_MT_*) are constant during a simulation run. The model points along the filament is spaced regularly at an interval of 0.1 *μ*m. The bending elastic modulus of the filament (*κ*), thermal energy, drag experienced by the filament, and the motor forces determine the filament dynamics with most values taken from literature (Table S1).

#### Motors

Motors are modeled as Hookean springs, represented as a bead with a spring (Figure 2(a)). The bead represents the motor head and the base of the spring depicts the tail which is immobilized on the surface. The motor head can bind to the MT with a probabilistic attachment rate of *r_a_* if it is within a certain distance to the filament, the binding range *d_a_*. The motors are modeled as continuous steppers with a fixed load-free velocity (*v*_0_) resulting in an instantaneous velocity *δ*v at each time step *δ*t, following a linear force-velocity relation:

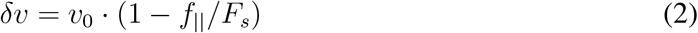

Here, the forces *f*_||_ and *F_s_* are respectively the component parallel the MT and the stall force. The motor is modeled to step forwards and backwards depending on the ratio of the parallel to stall force since the step size is given by: *δr* = *δv × δt* and *δv* can be positive or negative. The force experienced by the motors is calculated as *F_m_* = –*k_m_* · *δL_m_*, where *δL_m_* is the extension along the motor and the *k_m_* the stiffness constant of the spring. The motor can detach probabilistically with a detachment rate (*r_d_*), dependent on the magnitude of the force experienced by the motor under the stretch (*F_m_*). The detachment force (*F_d_*) is the characteristic detachment force of a single motor. Here, force-dependent detachment kinetics is modeled to be isotropic under assisting and hindering loads for simplicity, governed by the Kramers law (*34*):

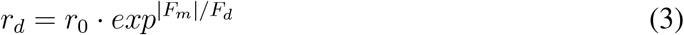

wherein, *r_d_* is the instantaneous and *r_o_* is the basal load free detachment rates of a single motor.

Motor parameters are also taken largely from experiment (Table S1).

### Materials and methods

#### Supporting methods

##### Protein purification and labelling

Tubulin was purified from goat brain using the high molarity PIPES based activity cycling protocol described for porcine brain tubulin (*35*) and used previously for goat brain tubulin (*23*). Tubulin was labelled with 5(6)-Carboxytetramethylrhodamine N-succinimidyl ester by incubating it with polymerized tubulin followed by a depolymerization-polymerization cycle to ensure activity of the labelled tubulin, as described in the EMBO Cell Biology Course 2003. Biotinylated and Alexa-488 labelled tubulin was prepared by the same method used for rhodamine labelling using either (+)-Biotin N-hydroxysuccinimide ester or Alexa Fluor 488 NHS-ester (Thermo Fisher Scientific, USA).

Yeast cytoplasmic minimal construct GST-Dyn331 (*20*) was expressed in yeast VY208 batch cultures induced with 2 % galactose(HiMedia, India) followed by affinity purification using the ZZ tag and pulled down using IgG beads (GE healthcare, Sweden). The protein was cleaved from the beads using a 6X-His tagged TEV which was expressed in E.coli BL21 DE3 and purified using Co-NTA resin (Thermo Scientific, IL, USA) followed by elution using imidazole (HiMedia, India).

In order to specifically immobilize dynein on glass surface, Anti-GFP nano-body was used. The pGEX6P1-GFP-Nanobody construct was a gift from Kazuhisa Nakayama (Addgene plasmid 61838 http://n2t.net/addgene:61838. RRID:Addgene_61838). The nanobody was expressed in E.coli BL21 DE3 by IPTG (SRL,India) induction at 1 mM for 4.5 hours at 30°C. The protein was purified from the clarified lysate using Glutathione Sepharose (GE health-care, Sweden) and eluted using reduced glutathione (*36*).

##### Gliding assay with pinned MTs

###### MT filament assembly

Plus end biotin labelled MTs were prepared by polymerizing 20 *μ*m unlabelled tubulin with 5 *μ*m rhodamine labelled tubulin in BRB80 and 10% glycerol at 37°C. The MTs were stabilized by addition of 20 *μ*m Taxol (Paclitaxel, Cytoskeleton Inc., CO, USA). Free monomers were removed by pelleting MTs at 150,000 g in TLA 100.3 rotor (Beckman Coulter, CA, USA). The MT pellet was resuspended in BRB-80 and 10% glycerol followed by addition of either 20 *μ*m biotin tubulin + 5 *μ*m rhodamine tubulin or 10 *μ*m each of Alexa Fluor-488 labelled tubulin and biotinylated tubulin. This mixture was incubated at 37°C for 30 minutes to continue polymerization of plus ends. Free monomers and aggregates were removed by centrifuging MTs at 130,000 g in TLA 100.3 rotor (Beckman Coulter, CA, USA) over a 60% glycerol cushion for 15 minutes. The MTs were re-suspended in BRB-80 and 20 *μ*m Taxol and used immediately.

The beating assays were performed in a flow-chambers constructed using 22 × 22 mm coverslip (No:1.5, VWR) adhered to a slide (22 mm x 60 mm, HiMedia, India) by two parallel strips of double backed Kapton polyimide tape (Ted Pella Inc., USA) spaced 7 mm apart, creating a chamber of area 22 mm x 7 mm.

###### Biotin-based plus-end tagging for streptavidin immobilization

The flow chamber was coated by 1:1 molar ratio mixture of nano-body and streptavidin. After blocking the surface with casein, dynein (1.18 *μ*g) was flowed in and allowed to incubate for 10 minute followed by plus end biotinylated MTs. The unbound dynein was removed by washing with Lysis Buffer (30 mM HEPES (HiMedia, India), 2 mM Mg-Acetate (Amresco, OH, USA), 50 mM K-Acetate (Fisher Scientific, India), 1 mM EGTA, 10% Glycerol). Plus end labelled MTs were flowed in and incubated for 10 mins. The unbound MTs were washed using wash buffer (Lysis Buffer + 20 *μ*M Taxol). The beating was observed after addition of motility buffer that consisted of lysis buffer supplemented with 4 mM ATP, Antifade mix (0.005 mg Glucose Oxidase, 0.0015 mg Catalase and 7.2 mM Glucose; all sourced from SRL Pvt Ltd., Mumbai, India). All assays were performed at 37°C. All reagents were from Sigma-Aldrich, MO, USA unless stated otherwise.

##### Microscopy and image Analysis

The filaments were imaged using a 60x Oil immersion lens (NA=1.45) mounted on a Nikon Ti-E inverted microscope (Nikon, Tokyo, Japan) using a TRITC filter with temperature control system (Okolab, Pozzuoli, Italy). The filaments were imaged every 10 seconds for 10-20 minutes using an Andor Clara2 CCD camera (Andor Technology, Belfast, UK). The images were denoised by applying median filter followed by background subtraction using ImageJ (*37*). Filaments were traced either interactively using NeuronJ plugin (*38*) or automatically using inhouse developed MATLAB code based on the ImageProcessing toolbox (Mathworks Inc., MA, USA). Time-projected images with encoding of time was based on the FIJI (*39*) plugin Zstack-DepthColorCode(https://github.com/ekatrukha/ZstackDepthColorCode). The XY coordinates of the tips were generated by tracking the tip using plugin MTrackJ 1.5.1 (*40*). Occasionally, some movies appeared to undergo systematic drift that was corrected by subtracting the XY coordinates of the stationary tip of MT from those of the moving tip. All analysis and plotting was performed in MATLAB (R2021b, Mathworks Inc., MA, USA). Dynein concentration per unit area (*μ*m^2^) was estimated as previously described (*23*) and briefly elaborated here. A range of concentrations of bacterially expressed and 6xHis affinity purified eGFP molecules were flowed into the flow chamber, imaged and molecules in the field of view were estimated by combining the image volume with the concentration. The image volume was estimated from the PSF and XY-area. The PSF was estimated by fitting a Gaussian profile to the fluorescence intensity of 0.1 *μ*m TetraSpeck beads (ThermoFisher Scientific, IL, USA) through the z-position (Figure S6(a)) with *PSF* ≈ 2.35*σ* (estimated by the parameter *d*). The scatter plot of the blank subtracted mean fluorescence intensity (y-axis) of increasing eGFP concentration in terms of area density in molecules per *μ*m?2 (x-axis) was fit to a straight line and the dynein-GFP mean intensity substituted. The resulting monomer density of 92.34 molecules/*μ*m^2^ resulted in a dimer density *ρ_m_* of 46.17 motors/*μ*m^2^ (Figure S6(d)).

##### Filament analysis

The filament movement is quantified from the coordinates of the free-end of the filament i.e. MT tip as displacement in x and y as a function of time. The span in 2D *s_x_* and *s_y_* is the difference between the maximum and minimum position occupied by the filament tip in the 2D space (*s_x/y_* = (*x/y*)_*max*_ – (*x/y*)_*min*_) (Figure S4). The elevation angle (*ϕ*) measures the angle subtended by the free-tip at the end of the pinning site with respect to its position at the time, t = 0 s. Another angular measure, the local tangent angle (*ψ*) is calculated at every discrete point after the end of the pinning site, along the filament contour as a measure of the shape of the filament.

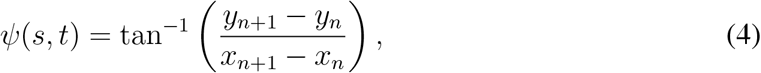

where (*x_n_*, *y_n_*) are the coordinates of the filament segment (*s_n_*) at time, *t*. The beat frequency, amplitude and the time period are inferred from time-series of the angle *ψ(s, t)*, that is smoothed using *smoothdata* (MATLAB, Mathworks Inc., MA, USA) and subjected to Fourier analysis. In experiments, filaments contours are obtained by manually tracking the microtubules using NeuronJ, an ImageJ plugin. All the analysis and plotting was performed in MATLAB (R2021b, Mathworks Inc., MA, USA).

The qualitative nature of the filament movement in 2D was quantified as the logarithm of the ratio of *s_y_* and *s_x_* as:

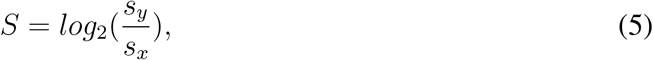

where *s_y_* = *y_max_* – *y_min_* and *s_x_* = *x_max_* – *x_min_* where *x_max_*, *y_max_* and *x_min_, y_min_* correspond to the maximal and minimal values of the position of the tip of the filament respectively.

##### Simulations

Simulations were run in a 2D space with a periodic boundary condition using an open-source agent-based C++ based Langevin dynamics environment Cytosim version f7921c4 from the git-lab (https://gitlab.com/f.nedelec/cytosim) (*41*). The effects of thermal energy and viscosity on the microtubule-motor interactions is also modeled, with k_*B*_T = 4.3 pN nm, viscosity *η* of 0.826 cP based on the viscosity of the buffer that contains 10% glycerol (*42*). Integration of simulations was done every 0.01 s.

### Supporting tables

**Table S1:** Parameters used in simulations. The values for mechanochemical properties of the MTs, motors and the simulation system are taken from literature where available.

| Symbol | Description | Value | Reference |
| --- | --- | --- | --- |
| *Microtubules* |  |  |  |
| $\kappa$ | MT bend-<br>ing rigidity | 20 pN $\mu\text{m}^2$ | (43) |
| $L_{MT}$ | MT length | 5-25 $\mu\text{m}$ | This study |
| *Dynein:* |  |  |  |
| $v_0$ | Single mo-<br>tor velocity | 0.10 $\mu\text{m/s}$ | (20) |
| $r_a$ | Attachment<br>rate | 5 1/s | (23) |
| $d_a$ | Attachment<br>range | 0.02 $\mu\text{m}$ | (23) |
| $F_d$ | Detachment<br>force | 3 pN | This study |
| $r_d$ | Detachment<br>rate | 0.04 1/s | (23) |
| $F_s$ | Stall force | 5 pN | (44) |
| $k_m$ | Linker<br>stiffness | 100 pN/$\mu\text{m}$ | (23) |

### Supporting figures

**Figure S1:**
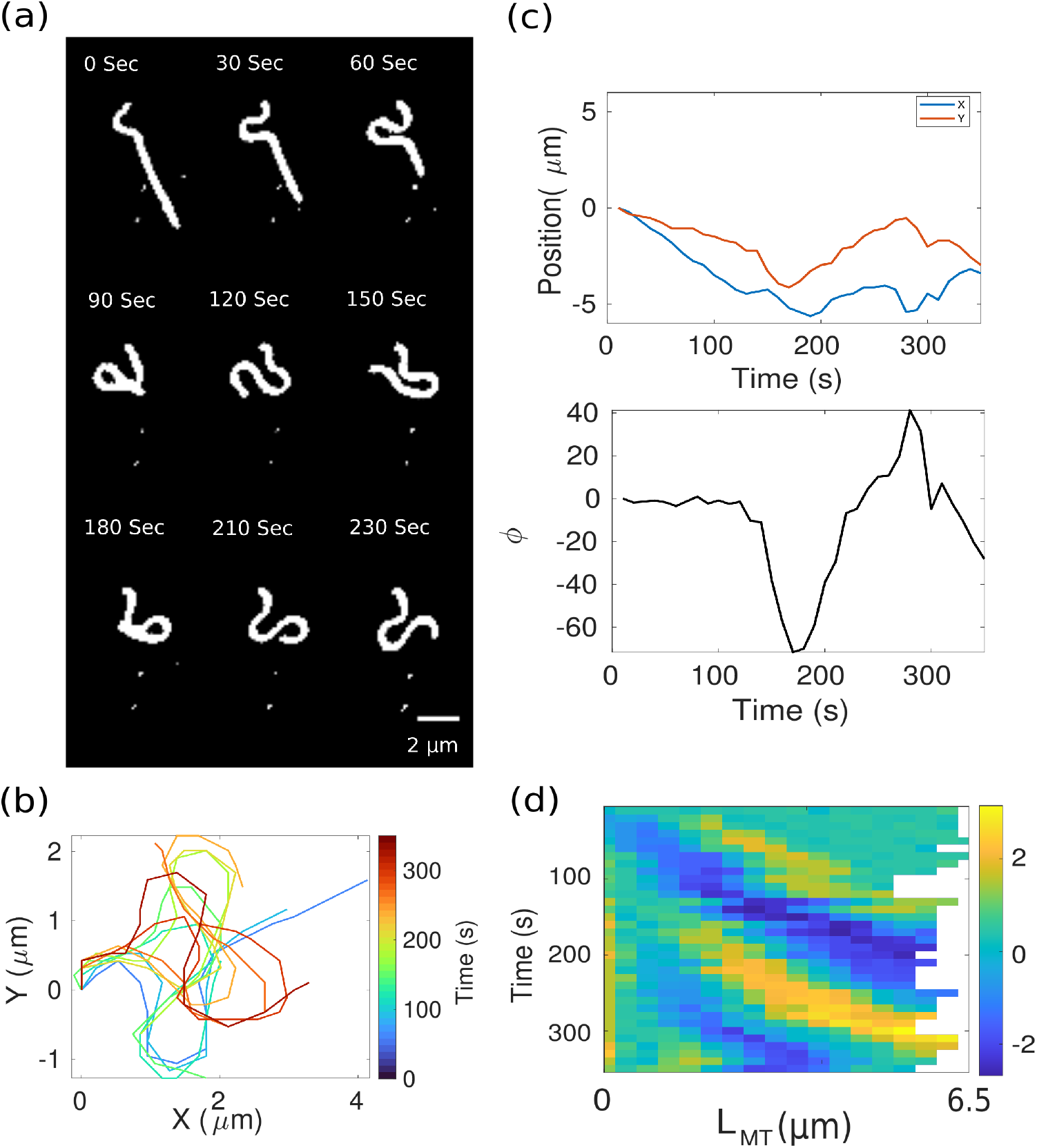
MT flagella-like waves of rhodamine-labelled MTs with plus-ends pinned. (a) Time-series of a representative MT labelled with rhodamine-tubulin demonstrating the emergence of flagella-like beating of a single plus end pinned MT, driven by surface-immobilized dynein (Video SV2). (b) Filament contours from experiment were projected in time at an interval of 30 s over the entire trajectory (color-bar: time in seconds). (c) The minus end of the MT that is free is analyzed for the position in X (blue) and Y (red) as a function of time (top), and its angular elevation *ϕ* (bottom) was plotted. (d) The local tangent angle along the MT contour, *ψ*, is plotted with time as a kymograph at 10 s intervals. Colors represent the angle in radians.

**Figure S2:**
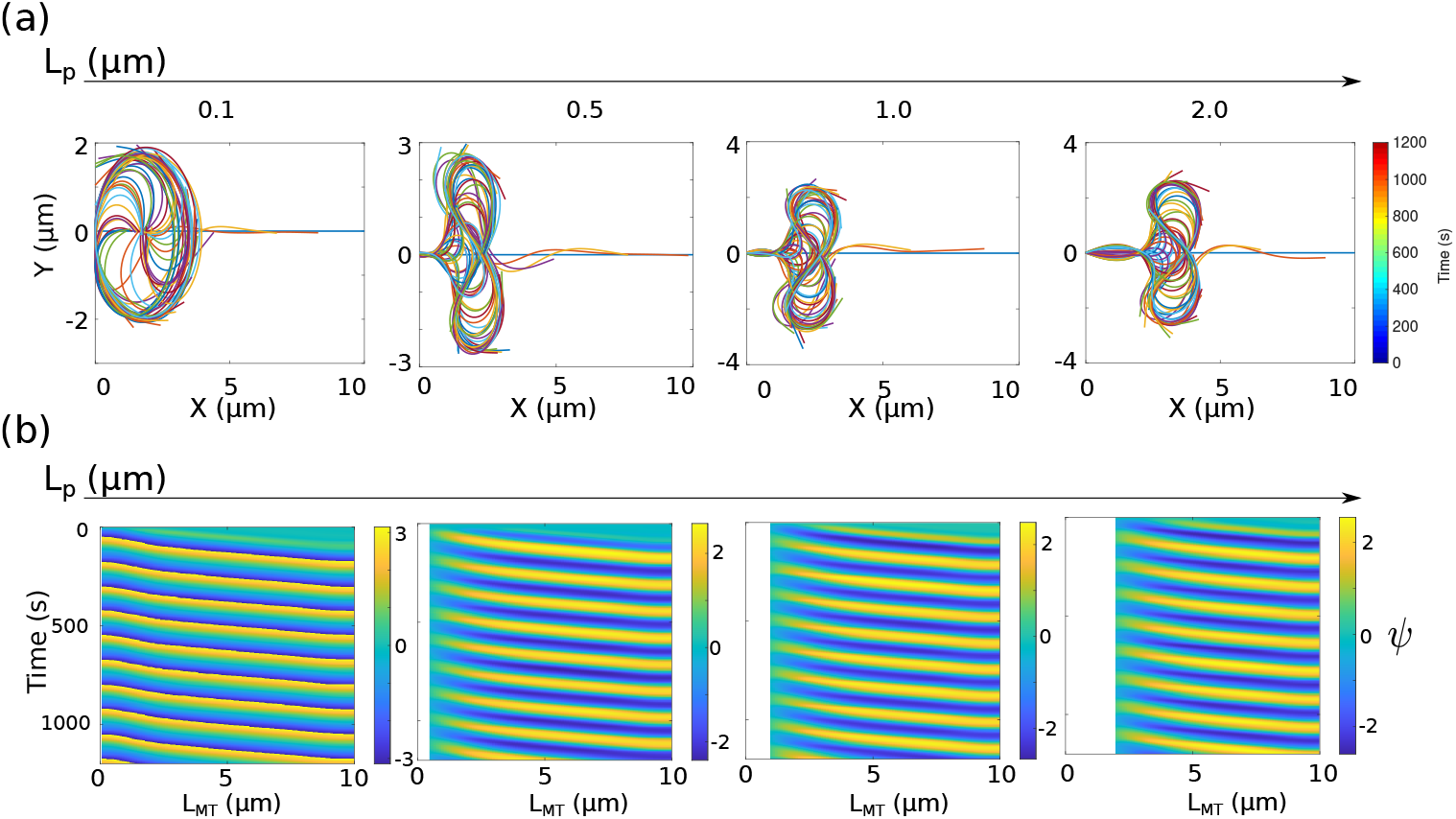
Effect of pinned length on patterns at a high motor density. The pinned MT length (*L_p_*) is varied for a constant length of the filament (10 *μ*m) and dynein density of 100 motors/*μ*m^2^. For increasing values of *L_p_*: (a) The time projected filament contours at every 30 s, and (b) kymographs of the tangent angles are represented.

**Figure S3:**
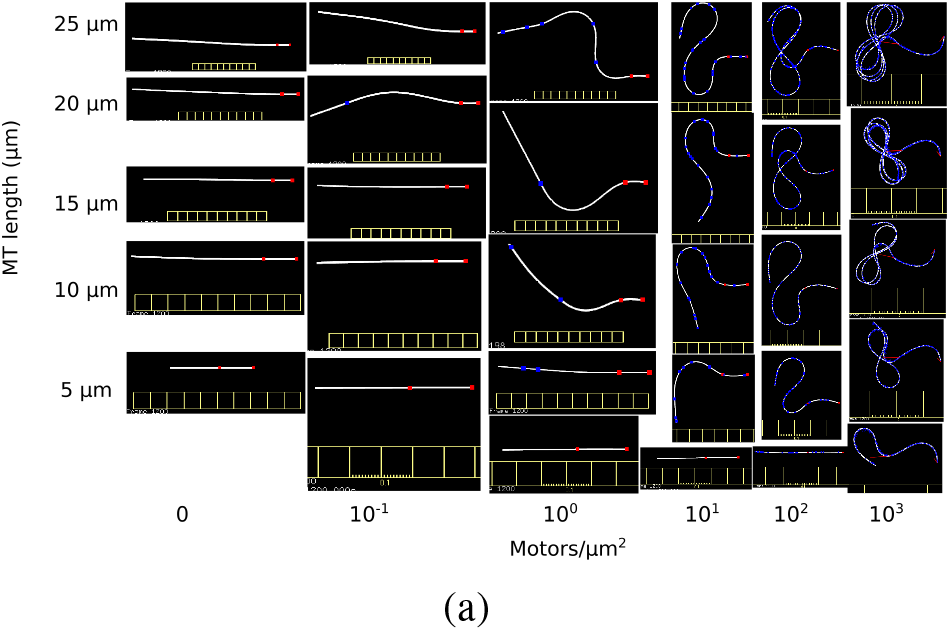
End-point traces of MTs simulated for increasing MT length and motor density. The snapshot of the simulation at the end of 1200 s for the varying lengths of MTs (along the rows) and densities (along the columns) with dynein motors (blue) is represented. The red marks indicates the pinning sites. Length of MT pinned is L_*p*_ = 2 *μ*m. Each vertical notch in the scale-bar: 1 *μ*m.

**Figure S4:**
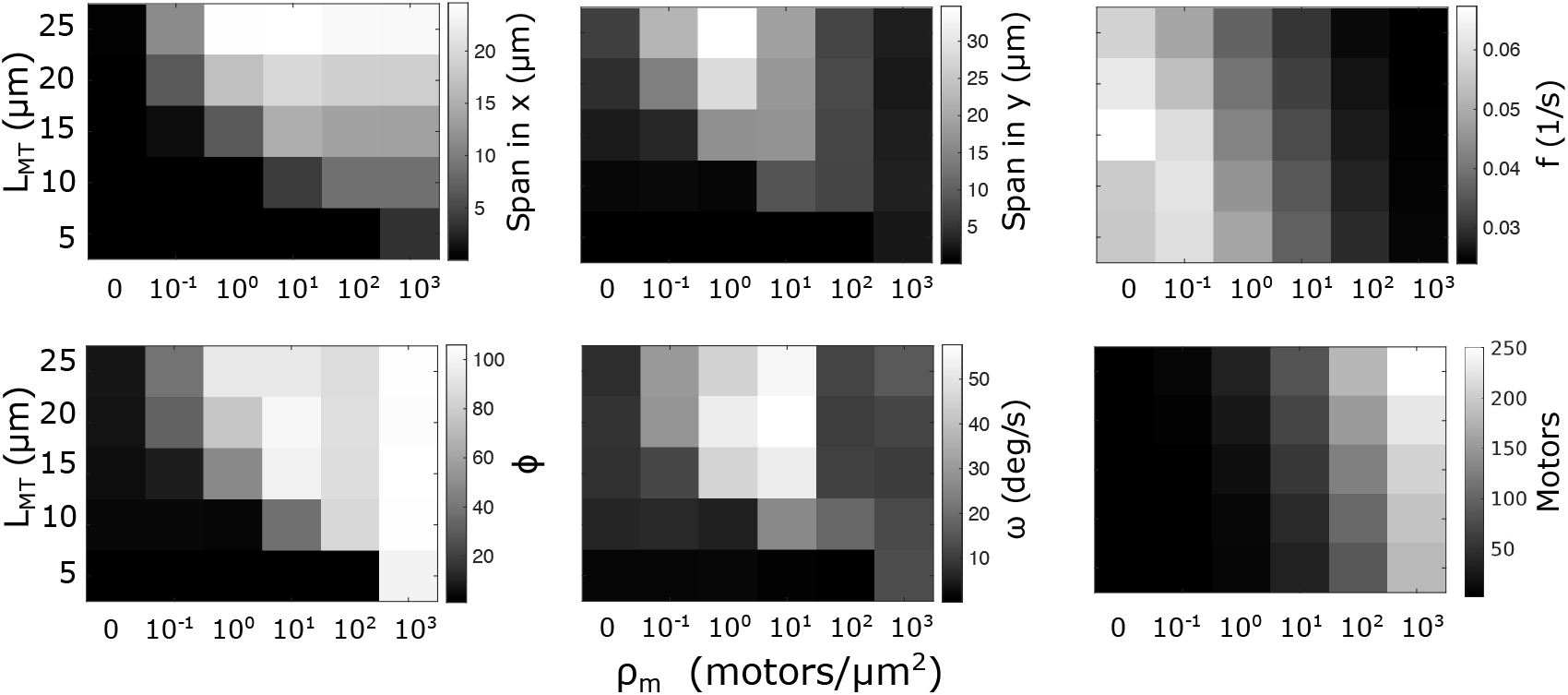
Quantifying filament movement dependence on motor density and MT length. The effect of varying motor density (*ρ_m_*) and MT lengths (*L_MT_* was quantified from multiple runs (*N_runs_*=10) from 600 s of simulations, when the system is at steady state. Mean values of the following measures are displayed: the span on the X- and Y-axis (maximal values, *μ*m), frequency (1/s) at the mid-point of the filament, the maximal elevation angle *ϕ*, maximal angular velocity of the tip *ω* and maximum number of motors bound to a filament (motors).

**Figure S5:**
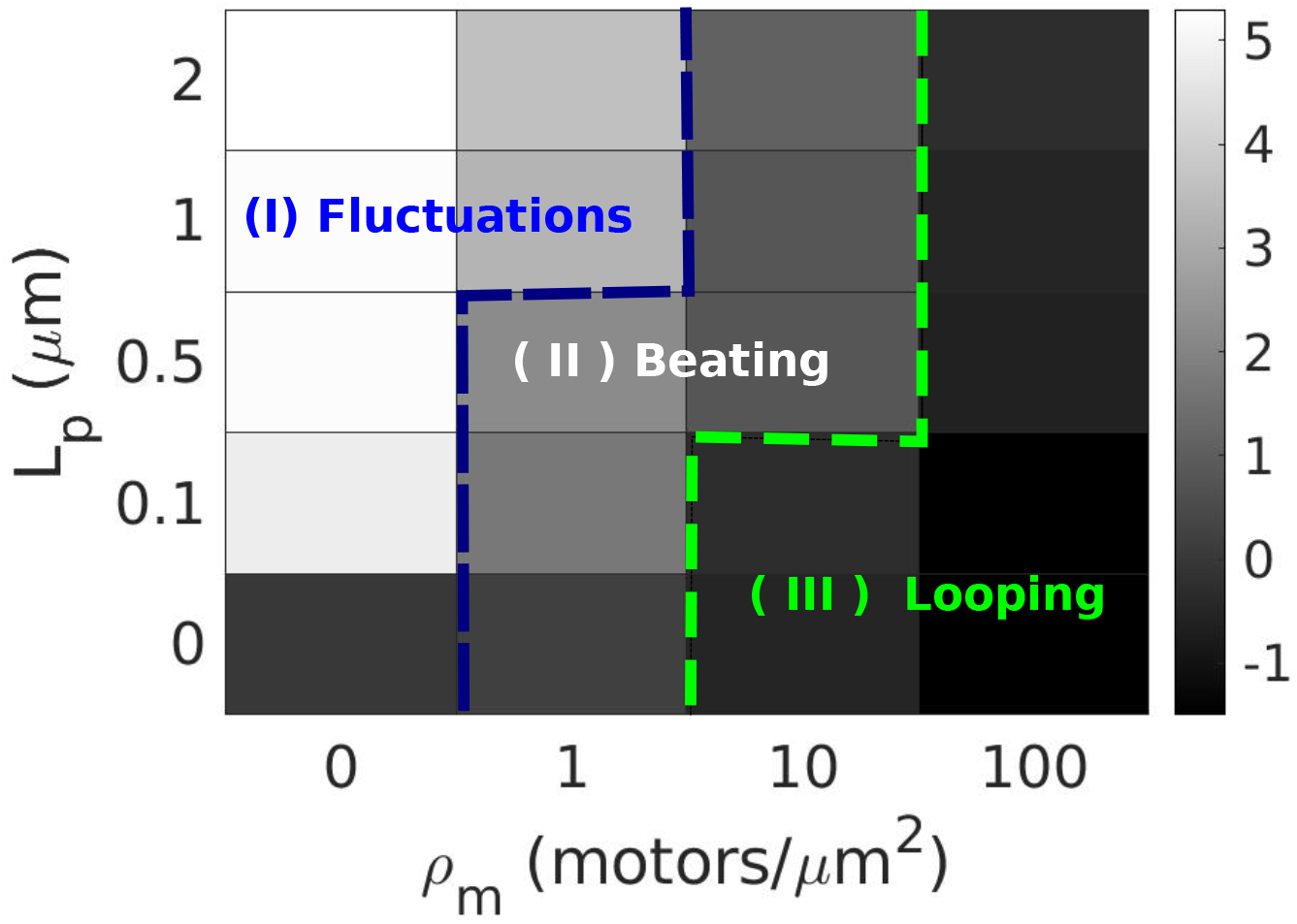
Simulating the phases of motion as a function of proportion of MT pinned and motor density. The parameter space for varying motor densities and %MT-pinned are divided into three distinct regions based on the derived measure, span (S) averaged over 10 independent runs. The three regions correspond to (I) fluctuations, (II) beating, and (III) looping. The MT length is constant, 10 *μ*m.

**Figure S6:**
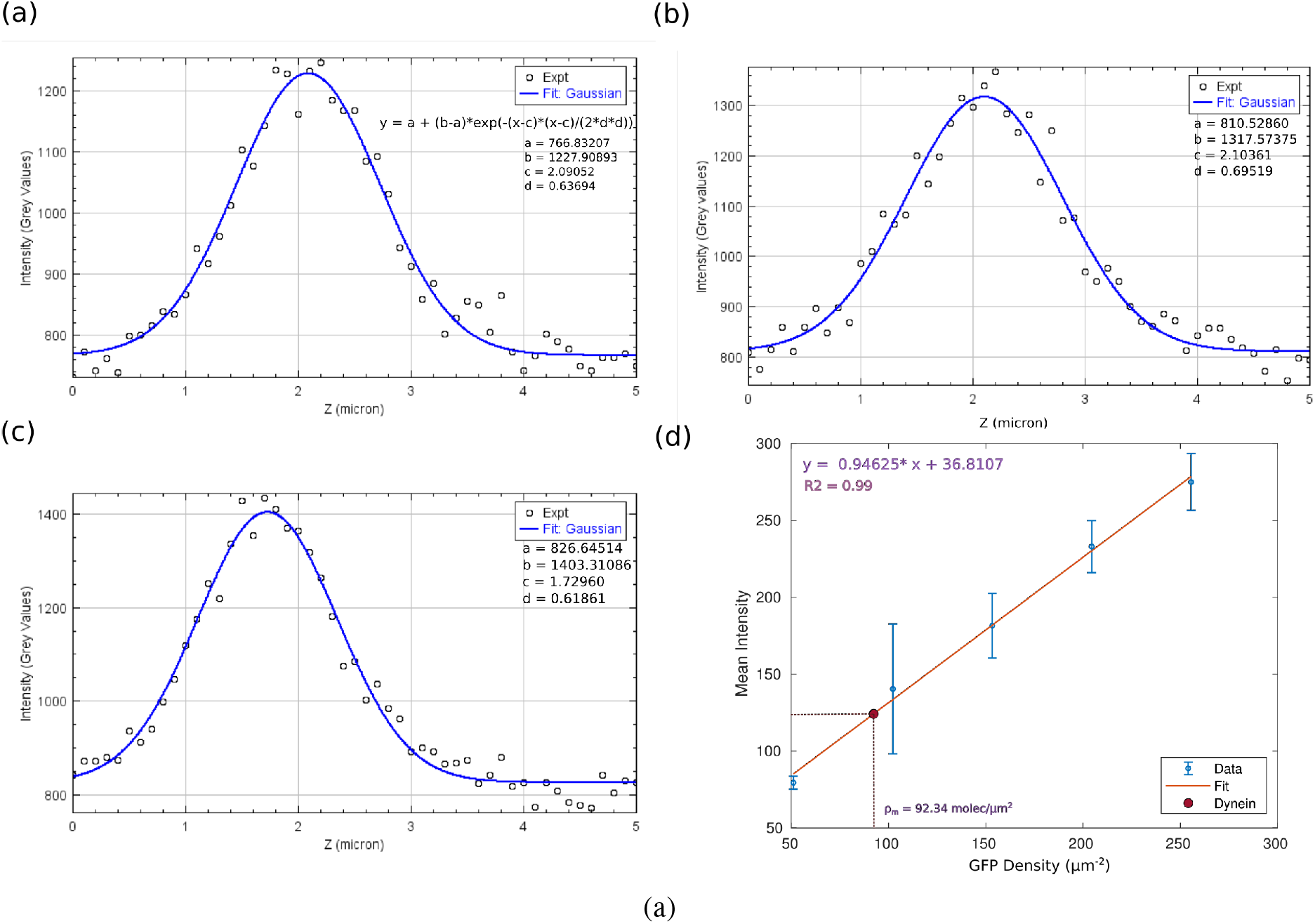
Dynein density estimation using EGFP standard calibration curve. (a) The PSF was estimated from the fluorescence intensity (y-axis) as a function of z-position (x-axis) of 0.1 *μ*m beads (red circles). The data was fit to a Gaussian (blue) described in the equation with the parameter d a measure of standard deviation. (b) A plot of fluorescence image intensity measured in FITC channel Vs number of molecules for a known standard range of EGFP with mean (blue) and standard deviation. The equation from the linear fit (Red) was used for calculating number of molecules in unknown i.e. dynein (Green) in the beating assay set up.

### 0.1 Supporting videos

**Video SV1:**
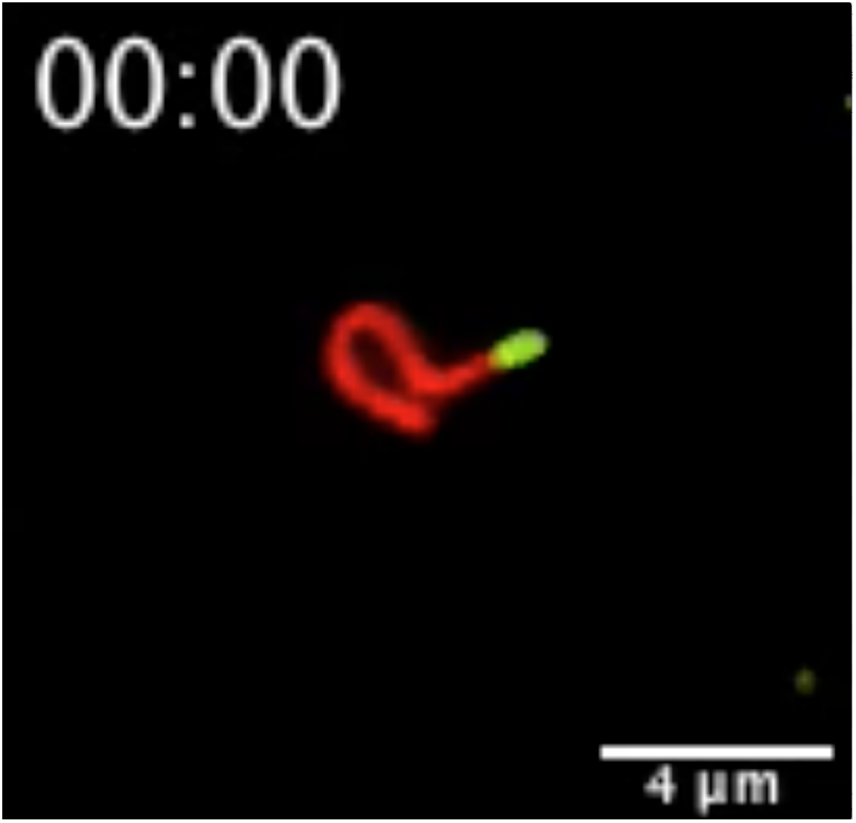
Flagella-like motion of plus-end pinned dual-color labelled MTs in a dynein gliding assay. A representative MT is seen in dual channel microscopy with an Alexa488 labelled plus end (green) and the minus end labelled with rhodamine labelled tubulin (red). The plus-end was co-polymerized with biotinylated tubulin and the coverslip was coated with a mixture of streptavidin and anti-GFP nanobody (1:1) molar ratio, passivated with casein followed by flowing in of a minimal GFP-dynein (GFP-Dyn) construct. The assembled MTs were flowed in and after incubation motor was activated by flowing in the motility buffer and imaged.

**Video SV2:**
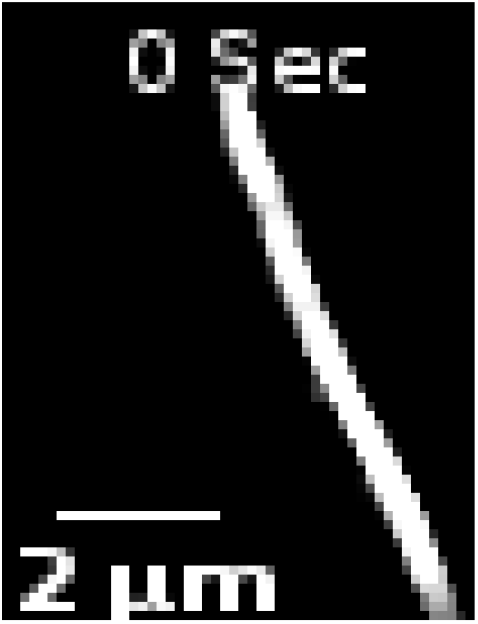
Single rhodamine labelled MT with biotinlyated plus-ends and free minus-ends. The movies represent magnified regions of data acquired with displaying single filaments that perform flagella-like beating in a dynein gliding assay. Pinning sites are provided by streptavidin coated surface with biotinylated tubulin labelling the plus-ends (*L_MT_* = 6.72 *μ*m, time= 340 s).

**Video SV3:**
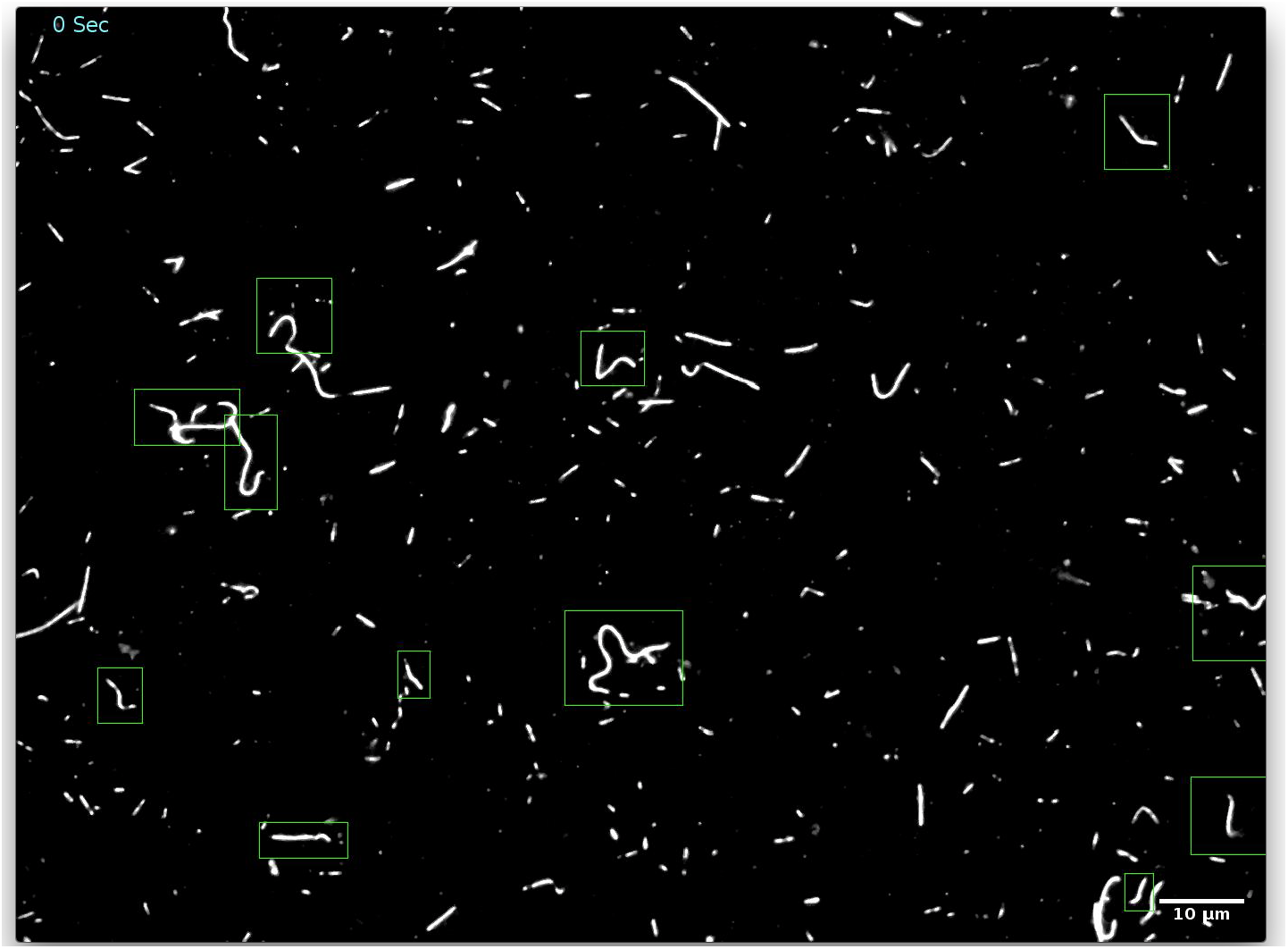
Dynein based flagella-like beating assay. The movies are whole fields of view of experimental time-series of fluorescently labelled MTs in a dynein driven flagellar-beating assay. The MTs are pinned to the surface by streptavidin coated surface with biotinylated tubulin labelling the plus-ends. Green boxes depict the location of filaments that appear to undergo flagella-like beating.

**Video SV4:**
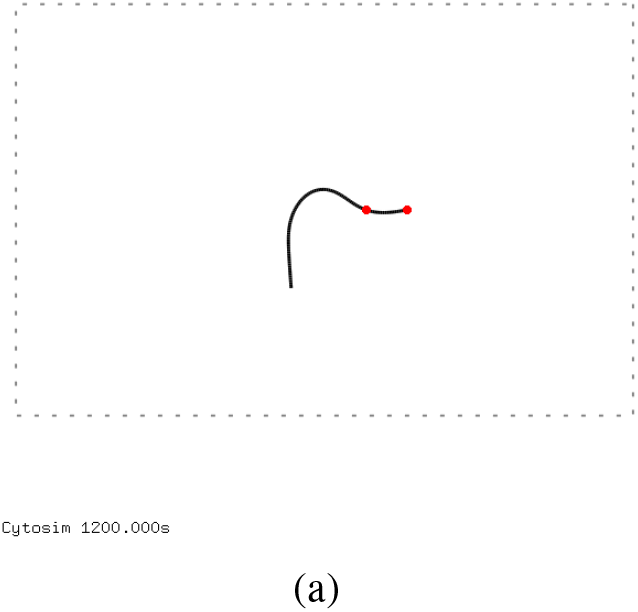
Simulating anchored MT beating driven by surface-immobilized motors. A simulation movie over the 330 seconds of a 10 *μ*m microtubule filament anchored at the plus end at two sites at a distance of 2 *μ*m (red). The motor density corresponds to 10 motors/*μ*m^2^.

**Video SV5:**
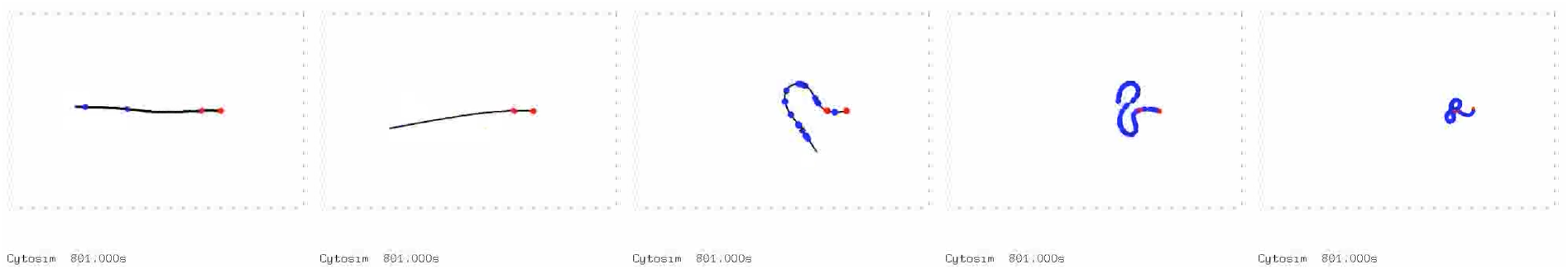
Simulating the effect of motor density on MT beating. Simulation movies in presence of increasing motor density, *ρ_m_* increases (from left to right) as 10^-1^, 10^0^, 10^1^, 10^2^ and 10^3^ motors/*μ*m^2^ with an MT of 10 *μ*m length with 2 *μ*m pinned to the surface at the plus-end. MTs are depicted as black lines, while motors bound to the MT are displayed as blue circles. Time: 20 min.

**Video SV6:**
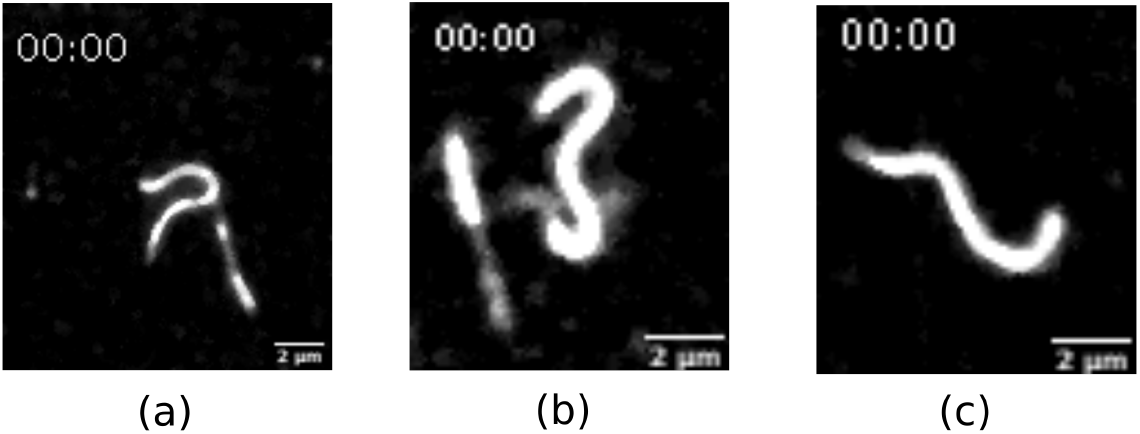
Effect of increasing pinning length on beating as observed in experiments. Time lapse movies of representative beating MTs with increasing proportion (a) 3.6%, (b) 6.4% and (c) 22.4% of the MT length pinned by biotinylated plus ends immobilized on a surface coated with streptavidin and cytoplasmic dynein. Time: mm:ss.

## Notes

### Competing Interest Statement

The authors have declared no competing interest.

## References

1. I. R. Gibbons, A. J. Rowe, Science 149, 424 (1965).

2. D. Nicastro, et al., Science 313, 944 (2006).

3. K. E. Summers, I. R. Gibbons, Proceedings of the National Academy of Sciences 68, 3092 (1971).

4. C. Gadelha, B. Wickstead, K. Gull, Cell Motility and the Cytoskeleton 64, 629 (2007).

5. G. Langousis, K. L. Hill, Nat Rev Microbiol 12, 505 (2014).

6. G. Cicconofri, G. Noselli, A. DeSimone, Elife 10 (2021).

7. I. H. RiedelKruse, A. Hilfinger, J. Howard, F. Jülicher, HFSP Journal 1, 192 (2007). PMID: 19404446.

8. P. Satir, T. Heuser, W. S. Sale, Bioscience 64, 1073 (2014).

9. M. E. Holwill, J. L. McGregor, J Exp Biol 65, 229 (1976).

10. P. Sugrue, M. R. Hirons, J. U. Adam, M. E. Holwill, Biol Cell 63, 127 (1988).

11. C. J. Brokaw, D. J. Luck, Cell Motil 3, 131 (1983).

12. C. B. Lindemann, Ann N Y Acad Sci 1101, 477 (2007).

13. C. B. Lindemann, K. A. Lesich, Cytoskeleton (Hoboken) 78, 36 (2021).

14. T. Okagaki, R. Kamiya, J Cell Biol 103, 1895 (1986).

15. L. Bourdieu, et al., Phys. Rev. Lett. 75, 176 (1995).

16. A. Vilfan, S. Subramani, E. Bodenschatz, R. Golestanian, I. Guido, Nano Lett 19, 3359 (2019).

17. T. Sanchez, D. T. N. Chen, S. J. DeCamp, M. Heymann, Z. Dogic, Nature 491, 431 (2012).

18. R. Sasaki, et al., Nanoscale 10, 6323 (2018).

19. I. Guido, et al., Active beating of a reconstituted synthetic minimal axoneme (2021).

20. S. L. Reck-Peterson, et al., Cell 126, 335 (2006).

21. S. Can, M. A. Dewitt, A. Yildiz, eLife 3 (2014).

22. M. P. Nicholas, et al., Proc Natl Acad Sci U S A 112, 6371 (2015).

23. K. Jain, N. Khetan, C. A. Athale, Soft Matter 15, 1571 (2019).

24. J. T. Canty, R. Tan, E. Kusakci, J. Fernandes, A. Yildiz, Annu Rev Biophys 50, 549 (2021).

25. F. Gittes, E. Meyhöfer, S. Baek, J. Howard, Biophys J 70, 418 (1996).

26. C. P. Brangwynne, et al., J Cell Biol 173, 733 (2006).

27. T. Sanchez, D. Welch, D. Nicastro, Z. Dogic, Science 333, 456 (2011).

28. K. Sekimoto, N. Mori, K. Tawada, Y. Y. Toyoshima, Phys. Rev. Lett. 75, 172 (1995).

29. L. Liu, E. Tüzel, J. L. Ross, Journal of Physics: Condensed Matter 23, 374104 (2011).

30. G. A. Monzon, L. Scharrel, L. Santen, S. Diez, J Cell Sci 132 (2018).

31. A. K. Rai, A. Rai, A. J. Ramaiya, R. Jha, R. Mallik, Cell 152, 172 (2013).

32. B. M. Friedrich, I. H. Riedel-Kruse, Elife 3, e03804 (2014).

33. M. Edamatsu, Biochem Biophys Res Commun 447, 596 (2014).

34. H. Kramers, Physica 7, 284 (1940).

35. M. Castoldi, A. V. Popov, Protein Expression and Purification 32, 83 (2003).

36. Y. Katoh, S. Nozaki, D. Hartanto, R. Miyano, K. Nakayama, Journal of Cell Science 128, 2351 (2015).

37. C. A. Schneider, W. S. Rasband, K. W. Eliceiri, Nature Methods 9, 671 (2012).

38. E. Meijering, et al., Cytometry 58A, 167 (2004).

39. J. Schindelin, et al., Nat Methods 9, 676 (2012).

40. E. Meijering, O. Dzyubachyk, I. Smal, Methods Enzymol 504, 183 (2012).

41. F. Nedelec, D. Foethke, New Journal of Physics 9, 427 (2007).

42. O. H. E. Segur J. B., Ind. Eng. Chem. 43, 2117 (1951).

43. F. Gittes, B. Mickey, J. Nettleton, J. Howard, J Cell Biol 120, 923 (1993).

44. A. Gennerich, A. P. Carter, S. L. Reck-Peterson, R. D. Vale, Cell 131, 952 (2007).

